# Computationally-efficient spatiotemporal correlation analysis super-resolves anomalous diffusion

**DOI:** 10.1101/2020.12.26.424447

**Authors:** Shawn Yoshida, William Schmid, Nam Vo, William Calabrase, Lydia Kisley

**Author notes:** Co-first authors with equal contribution (Shawn Yoshida, William Schmid).

## Abstract

Anomalous diffusion dynamics in confined nanoenvironments govern the macroscale properties and interactions of many biophysical and material systems. Currently, it is difficult to quantitatively link the nanoscale structure of porous media to anomalous diffusion within them. Fluorescence correlation spectroscopy super-resolution optical fluctuation imaging (fcsSOFI) has been shown to extract nanoscale structure and Brownian diffusion dynamics within gels, liquid crystals, and polymers, but has limitations which hinder its wider application to more diverse, biophysically-relevant datasets. Here, we parallelize the least-squares curve fitting step on a GPU improving computation times by up to a factor of 40, implement anomalous diffusion and two-component Brownian diffusion models, and make fcsSOFI more accessible by packaging it in a user-friendly GUI. We apply fcsSOFI to simulations of the protein fibrinogen diffusing in polyacrylamide of varying matrix densities and super-resolve locations where slower, anomalous diffusion occurs within smaller, confined pores. The improvements to fcsSOFI in speed, scope, and usability will allow for the wider adoption of super-resolution correlation analysis to diverse research topics.

## 1. Introduction

Anomalous diffusion in confined and crowded environments plays a role in the function of material and biophysical systems. In smart hydrogels, the anomalous diffusion of analytes within stimuli-responsive polymers are used for drug delivery and separation applications [1–4]. The heterogeneous environment of the extracellular matrix causes anomalous diffusion of signaling proteins, nucleic acids, or small molecule therapeutics that can affect the time scale of cellular changes [5]. Similarly, the topologic complexity of the intracellular space with cytoskeletal filaments and macromolecular crowding leads to anomalous diffusive behavior as biomolecules within the cell [6–8]. The effect of the heterogeneous nanostructures found in soft materials on the anomalous diffusion of molecules has important consequences in the use of biological, industrial, and medical materials.

Characterizing the interplay between the nanoscale structure of a material and anomalous diffusion is difficult with existing techniques. Scanning and transmission electron microscopy (SEM/TEM) require the sample to be prepared with flash-freezing or fixation that can warp the nanostructure of the materials, and cannot be used to quantify diffusion dynamics [9]. Atomic force microscopy (AFM) can also not quantify diffusion within materials and is limited to surface imaging only [10]. Optical techniques such as fluorescence correlation spectroscopy (FCS) and fluorescence recovery after photobleaching (FRAP) can accurately measure diffusion [11–17], but the diffraction-limited nature of light prevents the techniques from examining the heterogeneity of porous structures at nanoscales [18]. Super-resolution optical fluctuation imaging (SOFI) can super-resolve structure alone by analyzing wide field image stacks of independent, photoblinking, fixed fluorophores through the temporal correlation of pixel intensities and does not require sample preparation that might distort the material’s structure [19,20].

Correlation-based optical imaging can map diffusion at super-resolutions. fcsSOFI is an optical microscopy technique that extracts both super-resolution spatial and diffusion information. Introduced in 2015 by Kisley *et al*. [21], fcsSOFI uses a novel combination of two correlation-based techniques, FCS [22–25] and SOFI [19,20], to produce a super-resolved spatial map of diffusion dynamics within nanostructured materials. fcsSOFI analysis combines the two methods (Fig. 1). First, the intensity fluctuations of diffusing probe molecules (Fig. 1a) are recorded on individual pixels on a 2D camera (Fig. 1b) and correlated (Fig. 1c) at each pixel. SOFI is then used to convert the amplitude of the resulting correlation curves into a super-resolution image of the material nanostructure (Fig. 1d). FCS is then applied by fitting the correlation curves to a known diffusion model (Fig. 1e) [25] to extract diffusion information (Fig. 1f, h). Finally, spatial and diffusion information are combined with image fusion, which produces super-resolved maps of diffusion parameters (Fig. 1g, i).

**Fig. 1.**
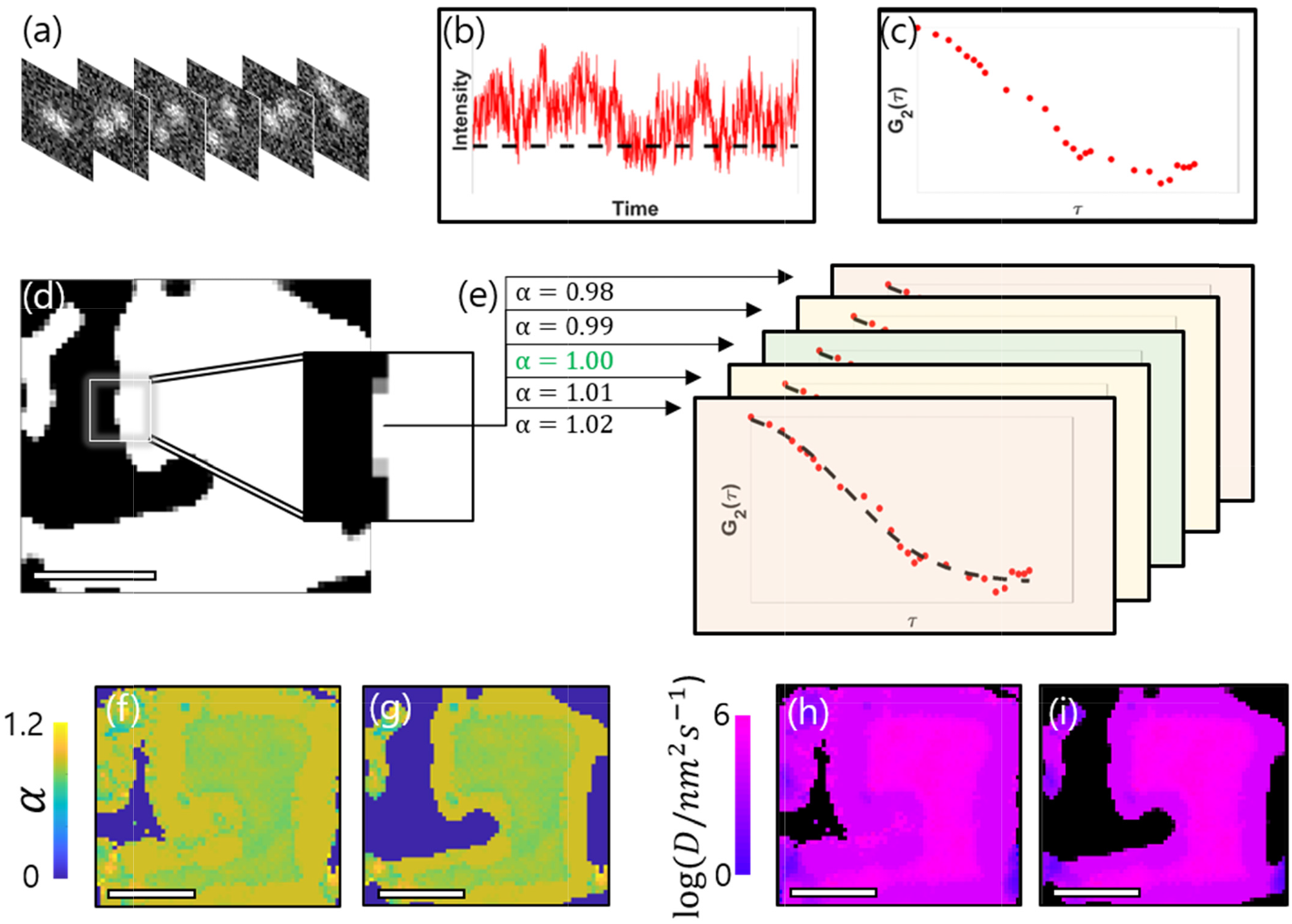
Steps of parallelized fcsSOFI applied to simulated anomalous diffusion. (a) Example frames of simulated anomalous diffusion of a fluorescent emitter in a porous material. (b) An intensity transient is extracted from each pixel, and (c) autocorrelation analysis is applied to each pixel’s transient. The value for G_2_(r,τ) at the first time lag, τ = 1, is used for the saturation of pixel r in the (d) SOFI image, where white represents a void, sampled area and black represents a barrier to diffusion. (e) Each autocorrelation curve is then fitted with the anomalous diffusion model using Gpufit, a parallelized iterative fitting toolkit [26]. At each pixel, multiple sets of fit parameters α (shown: true value, green; incorrect values, yellow, red) and D are tested simultaneously rather than iteratively. Color of plots in (e) are representative of the goodness of fit. The transition from serial to parallel computing decreases computation time. The extracted values of α and D are then used to create an (f) α-map and (g) D-map. They are each combined with the (d) SOFI image to create a super-resolved (h) α-map and (i) D-map. Scale bars are 1 μm.

fcsSOFI has been applied to a select set of materials by the developers. Previously, fcsSOFI was used to map diffusion in porous materials like agarose hydrogels and aqueous lyotropic liquid crystal gels [21], to extract diffusion information in confined, phase-separated polymers and biomolecules [27], and to relate the size and structure of a pH-responsive hydrogel to the interfacial transport properties of lysozyme proteins [1]. fcsSOFI was shown to extract diffusion coefficients more accurately and increase the spatial resolution of mapping pores more effectively at lower signal-to-background ratios than previous super-resolution imaging and single-particle tracking techniques. But the use of fcsSOFI analysis has been limited to these select applications by the developers Kisley and Landes, *et al*.

fcsSOFI has several limitations which hinder its use to more diverse datasets and by other research groups. fcsSOFI is most significantly limited by its computational expense. The least-squares curve fitting step requires overnight analysis times for typical microscopy images on a desktop computer. Ideally, fcsSOFI would be accessible without the need for high performance computing. Analysis of diffusion modes beyond simple Brownian diffusion, such as multi-component or anomalous diffusion, requires the explicit reprogramming of the FCS step: the fit model function and its parameters must be redefined, and the pixel-by-pixel least-squares curve fitting loop must be edited appropriately. In addition, the MATLAB analysis code is scattered in multiple scripts and functions and lacks user-friendliness.

Here, we have built and distributed an updated fcsSOFI software package which significantly improves upon the speed, scope, and accessibility of the original, while providing additional, biophysically-relevant diffusion models. First, following a widespread trend in computational and data science [28], we accelerate fcsSOFI by parallelizing the least-squares curve fitting step on a GPU. We then equip fcsSOFI with an expanded list of fitting models which allow for the analysis of experimentally-relevant modes of diffusion. We then apply fcsSOFI to simulated datasets of diffusing proteins in polyacrylamide images and demonstrate fcsSOFI’s ability to accurately characterize pore size and reveal non-Brownian diffusion dynamics due to confinement. Lastly, we package our updated software in a flexible, modular, and user-friendly graphical user interface (GUI), and distribute the accelerated code and GUI as an open-source tool via GitHub. With our updated fcsSOFI, we maintain previous levels of accuracy and performance while greatly decreasing runtimes, enabling the analysis of multiple datasets on standard computer hardware. We hope the accessible GUI enables a more diverse group of researchers to use fcsSOFI in their own work exploring diffusion within diverse samples of interest.

## 2. Computational Analysis and Improvements

### 2.1 Computational equipment

MATLAB, the language the original version of fcsSOFI was developed in by Kisley, et al. in 2015 [21], was used to implement the updates to the fcsSOFI software. MATLAB versions 2018a and 2020a were used to perform the updates. The open-source software build manager CMake (version 3.15.0) and Microsoft Visual Studio 2019 were used to incorporate GPU-parallelized curve fitting into fcsSOFI by rebuilding the MATLAB binding of the open-source curve fitting library Gpufit [26]. Execution speeds were benchmarked on two sets of computer hardware: a 2019 Dell Precision 3630 desktop equipped with a 3.7 GHz Intel Core i7 8th Gen CPU, an 8 GB Nvidia Quadro P4000 GPU, and 32 GB of RAM; and a 2017 Dell XPS 15 laptop equipped with a 2.8 GHz Intel Core i7 CPU, a 4 GB NVIDIA GeForce GTX 1050 GPU, and 16 GB of RAM.

### 2.2 GPU-parallelized, least squares curve fitting accelerates fcsSOFI computational time

Serial fitting is the most time-intensive step in the original fcsSOFI analysis. The iterative least-squares curve fitting in the FCS step is performed sequentially for each single pixel autocorrelation curve, where one set of parameter choices is evaluated before testing another set of parameters. Each single pixel autocorrelation curve can require many tens or hundreds of iterations to fit (an average of 86 fit iterations per pixel and maximum of 401 iterations were observed for a 30×30×5000 image stack). As a result, the FCS step can be a computationally demanding process, especially with experimentally-sized images, commonly 512 x 512 = 262,144 pixels or larger, with newer cameras containing upwards of 107 pixels [29].

Parallelization with GPU technologies can improve computation times in fcsSOFI. While modern CPUs are designed with compute cores that are, on their own, significantly faster than individual GPU cores, it is uncommon for a CPU to have more than eight cores (our laptop and desktop CPUs contain four and six cores, respectively), whereas commercial GPUs can consist of well over one thousand cores (the Nvidia Quadro P4000 contains 1792 cores). GPU-parallelization is thus most appropriate for tasks that require many small, but mutually independent computations, like iterative fitting. GPU computing has seen wider adoption in computational microscopy as parallelization can accelerate many of the demanding image processing tasks necessary to resolve large amounts of multidimensional microscope data [30]. GPU computing has previously been used to accelerate maximum likelihood estimation and localization-based super-resolution microscopy [31] as well as medically-relevant scanning photoacoustic microscopy [32].

We parallelize the pixel-by-pixel least-squares curve fitting step that extracts diffusion information in fcsSOFI with an open-source GPU-based curve fitting method. Gpufit is an open-source, CUDA-based GPU curve fitting package developed by Przybylski *et al*. in 2017 [26]. The developers of Gpufit demonstrated the efficacy of their GPU curve fitting package with an application to a stochastic optical reconstruction microscopy (STORM) algorithm [26]. Here, Gpufit mediates the computational expense of the original FCS implementation by parallelizing the iterative fitting on a GPU. Since Gpufit was developed for fitting and localizing single-molecule diffraction-limited patterns, the original binding does not include the fit model functions for the various modes of nanoscale diffusion we study. We used the functionality of Gpufit for user-defined custom models in the form of CUDA header files and corresponding model IDs via C programming [26]. Implementing Gpufit maintains the same accuracy and precision as the original MATLAB ‘fit’ function in the original CPU-based fcsSOFI. In the case of fcsSOFI, instead of performing each fitting iteration sequentially on a CPU, the now parallelized FCS step tests many parameter combinations simultaneously by placing each independent fit iteration on one of the many hundreds of compute cores on a GPU (Fig. 1e).

fcsSOFI using GPU-parallelized fitting can be upwards of 40 times faster than CPU fitting depending on image size. The execution time of the updated, parallelized and original, serial fcsSOFI codes were studied as data size was increased (Fig. 2). An experimental set of 100 nm fluorescent beads-diffusing in agarose hydrogel data used in the original study on fcsSOFI [21] was segmented into five region-of-interest (ROI) sizes ranging from 32×32 pixels to the original size of 512×512 pixels. All 1000 image frames from the original data set were used for each ROI size, and all five constructed data sets were fed into both GPU- and CPU-based fitting versions of fcsSOFI. Execution times were recorded using the ‘tic’ and ‘toc’ functions in MATLAB. For the GPU-parallelized code, three separate trials were taken at each ROI size and averaged. The Dell Precision desktop computer was used for every trial. The updated code significantly outperformed the original code at every image size due to GPU-parallelized fitting: approximately 40 times faster for the 32×32-pixel image and still 10 times faster for the experimentally sized 512×512-pixel image (Fig. 2). In particular, the 32×32-pixel image required 315.8 s to analyze with the original CPU-based fcsSOFI, an execution time which GPU-parallelized fitting improved to 7.8 ± 0.2 s, on average.

**Fig. 2.**
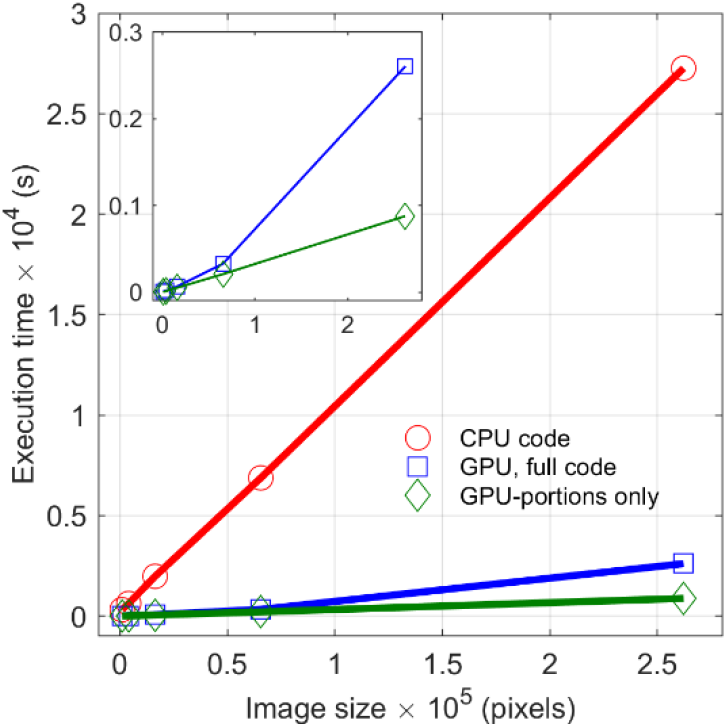
Parallelized, GPU fitting (blue) speeds fcsSOFI up to 40-times faster than sequential, CPU fitting (red). Total fcsSOFI execution times were measured at five experimental datasets ranging from 32 × 32 pixels to 512 ×512 pixels, 1000 frames long. Error bars (standard deviation) were between 0.05 and 2.5% of the average computation times and are too small to be viewed on the plot. The updated execution times are compared to the execution times of the parts of the code parallelized with Gpufit (green) and are magnified in the inset. Whereas the portions of code which are GPU-specific show a linear increase in computation time for larger image sizes, the entire updated code shows a disproportionate increase.

Despite the speed increases brought by the GPU-parallelization, the execution time improvement for larger images is limited by the proportion of the code that is not able to be parallelized and must be executed serially by the CPU. Whereas the execution times for the CPU-based code increase linearly as image size increases, which is expected for a proportional increase in computations for larger image sizes, Fig. 2 reveals a nonlinear increase in execution time for the GPU-based code. While this nonlinear trend initially looks unusual, we can look to Amdahl’s law for an explanation, which states that the maximum speed improvement *S* resulting from the GPU-parallelization of a given program is constrained by

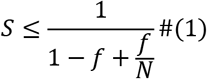

where *f* is the fraction of code that can be parallelized and *N* is the number of GPU cores [33]. Only tasks which consist of many small, independent computations are able to be executed in parallel, which is not the case for the many steps in the fcsSOFI algorithm, such as loading data into the MATLAB workspace, the SOFI analysis, creating figures, and saving data. Thus, the speed increase of the updated fcsSOFI software is limited by its large portion of code that is serially computed on the CPU (Fig 2., inset). Nevertheless, the GPU-parallelized code is still significantly faster than the original code at every image size tested.

### 2.3 Additional curve fitting models expand the relevance of fcsSOFI to other diffusion types

An expanded library of curve fitting models in fcsSOFI accounts for different types of diffusion. FCS is performed pixel-by-pixel by treating each pixel as an individual focal volume. The autocorrelation curve of each single pixel intensity trajectory is fit to a known model of diffusion [21]. In the original fcsSOFI code, autocorrelation curves were fit exclusively to a Brownian diffusion model [34], given by

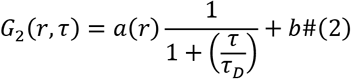

Here *G*_2_(*r, τ*), the autocorrelation for a position (pixel) *r* and time lag τ, is expressed in terms of an amplitude *α*(*r*). an offset *b*, and a characteristic diffusion time *τ_D_*. In our updated fcsSOFI software, the Brownian model given by Eq. 2 can be swapped with additional diffusion models without explicit reprogramming. The anomalous diffusion model is of particular relevance to diffusion in complex, crowded environments [35,36], and is given by

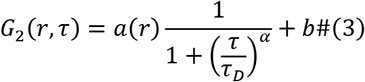

where *α* is the anomalous power [1]. The anomalous model reduces to the Brownian model for *α* = 1, where values of *α* < 1 indicate sub-diffusive dynamics, and *α* > 1 indicate super-diffusive dynamics. Sub-diffusive dynamics are common in crowded environments such as in cells, and confined environments like biological matrices, where motion of molecules may be impeded by particles and nanostructures within the environment. This results in a nonlinear relationship between the mean squared displacement (MSD) and time. Simulations demonstrate *α* can be accurately extracted with Eqn. 3 (Appendix A).

A two-component Brownian diffusion model is also available, given by

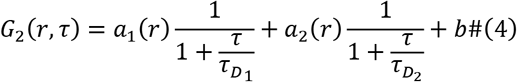

where the amplitude and characteristic diffusion time for each component is labeled with a 1 or 2.

Maps of the diffusion coefficient and anomalous power are produced with the curve fits at each pixel. The diffusion coefficient *D* is extracted for each fit from our two-dimensional microscopy data with

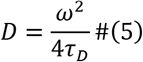

where *ω* is the size of the detection region and *τ_D_* is the best fit estimate of the characteristic diffusion time [37]. For fcsSOFI, *ω* can be taken as the physical width of each pixel, usually on the order of tens of nanometers. The values of *D* for each pixel are then compiled into a single *D* map where color scaling indicates the value of *D* across the image (Fig. 1i). For the exception of the expanded list of diffusion models used to extract an estimate for *τ_D_*, this step is largely unaltered from the original algorithm. However, with the addition of the anomalous diffusion model to updated fcsSOFI pipeline (Eq. 3), similar color maps can be made for the anomalous power *α* (Fig. 1g). The updated fcsSOFI software automatically produces the *α* map along with the FCS *D* map when the anomalous model is selected. If the two-component Brownian diffusion model is selected instead, a second map is produced for the second diffusion constant.

## 3. Simulation of biophysical protein diffusion under confined conditions

We produce and analyze simulated data of protein diffusion within hydrogel environments to demonstrate the applicability of the new functionalities in fcsSOFI to biophysically-relevant diffusion in nanostructure materials. Hydrogels are used to mimic the extracellular matrix as supports in tissue engineering and are used as drug delivery vectors [38,39]. From the perspective of proteins, the hydrogel presents local, nanoscale, physicochemical variations in steric confinement and chemistry that can affect the adsorption, diffusion, conformation, folding, and aggregation of the protein [40,41]. The in situ, super-resolution, quantitative capabilities fcsSOFI make it uniquely suited to produce images at the length-scales of hydrogels and the extracellular matrix and the time-scales of protein dynamics.

Environments with clearly segregated void space and gel barriers to simulate diffusion within were created by binarizing and morphologically altering existing SEM images of polyacrylamide from Lira *et al*. [42]. To account for uneven illumination in the SEM image, Otsu’s method, an adaptive algorithm, was used to binarize the image. Combinations of the built-in MATLAB morphological functions, ‘imclose,’ ‘imopen,’ ‘imdilate,’ and ‘imerode,’ were applied to the binarized image to mimic denser and more open polyacrylamide gels that emulate extracellular matrix from different areas of the body [43]. These morphologically altered binarizations (dense and open matrices), along with an original, unaltered binarization (medium matrix), were used as the environments for diffusion (Fig. 3a-c).

**Fig. 3.**
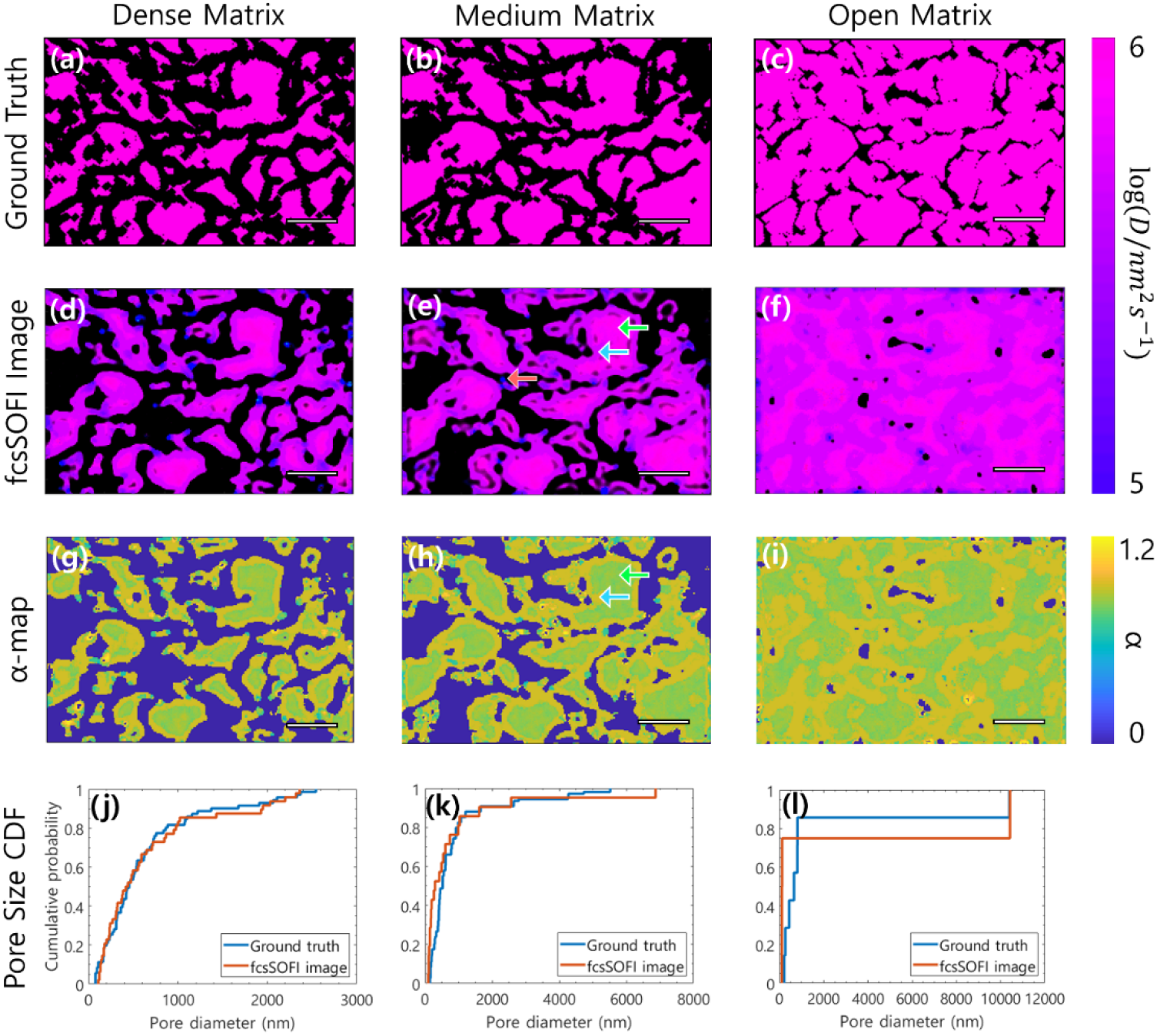
fcsSOFI extracts super-resolved maps of the diffusion dynamics of simulated fibrinogen diffusing in three densities of polyacrylamide gels. Ground truth images (a-c) were generated by coloring the binarized image used in the simulations with the color corresponding to the simulated diffusion coefficient, *D* = 1 × 10^6^ *nm*^2^ *s*^−1^. Super-resolved maps of diffusion coefficient (d-f) and anomaleity (g-i) were extracted by fcsSOFI. We observe the effects of confinement (e, h: green and blue arrows) and trapped diffusion (e: red arrow). Cumulative distribution of pore sizes for each matrix density verify fcsSOFI’s ability to resolve individual nanopores and produce accurate pore size distributions. Scale bars are 2 μm.

**Fig. 4.**
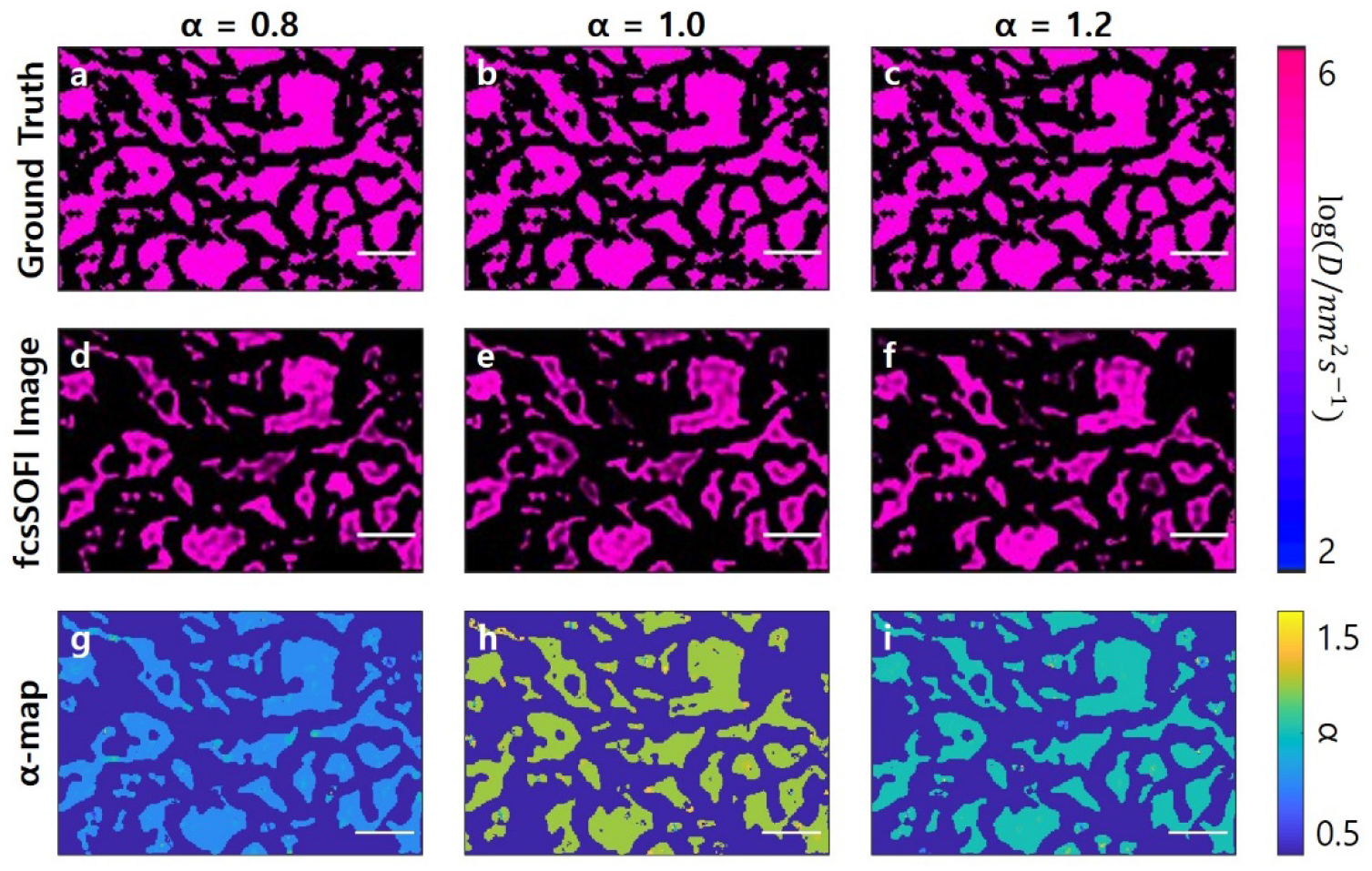
Comparison of the ground truth super-resolution diffusion map (a-c) and diffusion maps extracted by fcsSOFI (d-f) from three datasets with varying anomaleities. (a,d,g) are from a dataset simulated with *α =* 0.8, (b,e,h) with *α* = 1.0, and (c,f,i) with *α* = 1.2. All datasets were simulated with *D* = 1 × 10^5^ *nm*^2^*s*^−1^. The ensemble-averaged D values in (d,e,f) agreed within error, with *D*_*α*=0.8_ = (1.0 ± 0.4) × 10^5^ *nm*^2^*s*^−1^, *D*_*α*=1.0_ = (1.0 ± 0.4) × 10^5^ *nm*^2^*s*^−1^, and *D*_*α*=1.2_ = (1.0 ± 0.6) × 10^5^ *nm*^2^*s*^−1^. The ensemble-averaged *α* values in (g,h,i) also agreed with simulated value, with *α*_*α*=0.8_ = 0.81 ± 0.03, *α*_*α*=1.0_ = 1.0 *±* 0.1, and *α*_*α*=1.2_ = 1.20 ± 0.06. Scale bars are 2 μm.

**Fig. 5.**
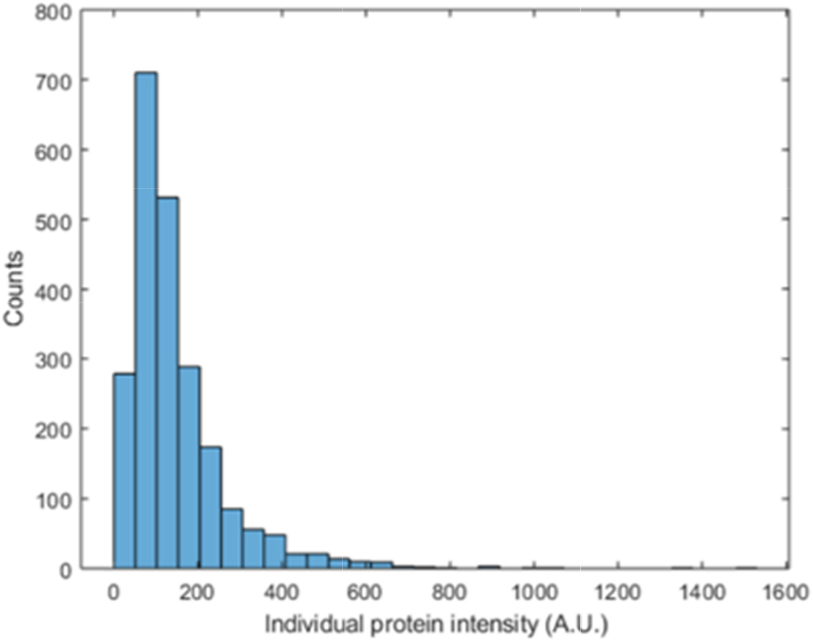
Distributions of individual protein intensities for 2260 fibrinogen molecules taken from the first frames of three movies collected at 10 mW laser intensity and 50 ms camera integration time. For fibrinogen, the average protein intensity with the local background removed is 150 ± 120 counts and the average local background is 140 ± 20 counts.

To simulate biophysically-relevant diffusion, 2D random walks were simulated within the nanocavities of the pore maps using step size distributions based on experimentally-obtained parameters (Appendix B). Fluorescently-labeled fibrinogen immobilized on glass was measured experimentally on a wide-field total internal reflection fluorescence (TIRF) microscope (Appendix C). The experimental values of the protein intensity and background were used in the simulation. Emitters were represented by 2D Gaussian point spread functions with a full width half max of 302 nm, and were simulated diffusing at *D* = 1 × 10^6^ *nm*^2^*s*^−1^ [44]. Each emitter’s intensity was drawn from a Poisson distribution to simulate shot noise, while simulated readout noise was added using a random normal distribution. An experimentally-determined signal to background ratio of 1.07 was used. Simulations were performed with a frame rate of 100 Hz and a pixel size of 47.6 nm [42]. Each step of the random walk was calculated based on the diffusion coefficient, time step, pixel size, and distribution width. It was then segmented into 1000 smaller substeps, and added onto the previous position substep by substep until the particle collided with a barrier or completed the full step. Particles colliding with barriers had their step sizes truncated at the collision point. Simulations of the Brownian diffusion of 300 emitters were recorded for 5000 frames and analyzed with fcsSOFI.

Super-resolved diffusion maps extracted by fcsSOFI show the local diffusion coefficients of the simulated emitters at the nanoscale (Fig. 3d-f). The ensemble averaged diffusion coefficient in the medium matrix is *D* = (5 ± 1) × 10^s^ *nm*^2^*s*^−1^ compared to the simulated *D* = 1 × 10^6^ *nm*^2^s^−1^. Examining local diffusion behavior below the diffraction limit reveals that while diffusive behavior in the centers of larger pores is around *D* = 1 × 10^6^ *nm*^2^*s*^−1^ (Fig. 3e, green arrow), diffusion coefficients near the edge of the pores, where emitters have a chance to move in the direction of the wall and immediately collide with the barrier, effectively lower the diffusion coefficient to around *D* = 5 × 10^s^ *nm*^2^*s*^−1^ (Fig. 3e, blue arrow). Furthermore, in the smaller pores and other more confined regions, emitters become trapped, with diffusion coefficients around *D* = 1 × 10^4^ *nm*^2^*s*^−1^ (Fig. 3e, red arrow). Trapped particles with lower diffusion coefficients were observed more frequently in denser matrices. The denser the matrix, the more pores there are with sizes comparable to the diameter of emitters. This results in some particles being trapped in pores that heavily restrict their step sizes, hence very low diffusion coefficients.

Anomalous factor maps (α-maps) extracted by fcsSOFI show the sub-diffraction limit effects of confinement on the local anomaleity of the simulated emitters. The ensemble averaged anomaleity, α, in the unaltered binarized SEM image was *α* = 0.87 ± 0.06 compared to the *α* = 1 expected of Brownian diffusion. Near the edges of larger pores, anomaleities around *α* = 1 (Fig. 3f, blue arrow) are observed as once a particle moves in a direction away from a barrier, they are likely to be able to take a full step. In the centers of pores, anomaleities < 1 (Fig. 3f, green arrow) are observed as step lengths are more likely to be cut short due to barriers being nearer to the emitter in every direction. The heterogeneities in both the diffusion coefficients and anomaleities appear in features below the diffraction limit, emphasizing the importance of the super-resolution capabilities of fcsSOFI.

Accurate pore size distributions were extracted by fcsSOFI. The super-resolution images (Fig. 3e-f) were compared to the ground-truth binarized and morphologically-altered SEM images used as the diffusion simulation environments (Fig. 3a-c). Pore sizes were extracted from both sets of images using Delauney triangulation [45] and cumulative distribution functions (CDFs) were generated to compare the pore size distributions (Fig. 3j-l). The overlap in the simulated and ground-truth CDFs is high, demonstrating the accuracy of fcsSOFI for spatial analysis. Notably, some isolated pores are missing from the analyzed image in cases where a pore was not sampled by a fibronogen molecule over the course of the simulation. This is likely to occur experimentally, not just in simulation, as fluorophores introduced to a porous medium have little chance of entering completely isolated pores.

## 4. Development of open-source fcsSOFI GUI

With the goal of making fcsSOFI accessible to wide array of researchers, we present our updated fcsSOFI code both as an easy-to-use GUI and modular MATLAB script. Accessible via MATLAB or the MATLAB compiler, the GUI enables fellow researchers to access the full functionality of fcsSOFI without having to interface with the source code or conduct any independent programming. The GUI contains clearly labeled text inputs, switches, and buttons for data input, diffusion model selection, data inspection, post-processing, figure modification, and exporting data. The GPU-parallelized fcsSOFI software is also available as a standalone MATLAB script for experienced programmers to have finer control on the analysis pipeline. Both the script and GUI automatically pair with the same custom Gpufit library containing the additional diffusion models, and are written to have as similar functionality as possible to accommodate varying preference and experience.

The fcsSOFI GUI, script, and customized Gpufit binding are openly distributed via GitHub [46]. Included in the GitHub distribution is a PDF user-guide for the GUI containing thorough user instructions and troubleshooting tips, as well as many orienting screenshots and diagrams. Researchers with interests in nanoscale diffusion and structure, super-resolution microscopy, and single particle tracking are able to freely download and use fcsSOFI on their own projects. While we anticipate our GUI to make fcsSOFI more accessible to researchers with limited coding skills, researchers who program frequently will be able to provide valuable feedback, feedback which can inform future updates to fcsSOFI. In addition, the fcsSOFI GUI and script will continue to be maintained by our lab with Git version control and updates will be pushed the public GitHub repository as the updated fcsSOFI software is further refined.

## 5. Conclusions

The nanoscale environments of porous, biophysical and material systems have macroscale implications on diffusion dynamics within them. We introduced an updated version of fcsSOFI, a correlation-based super-resolution technique that has previously been shown to be able to extract nanoscale structure and Brownian diffusion dynamics with improved speed, scope, and usability. The speed was improved by parallelizing the least-squares curve fitting step on a GPU, and achieve improved computation times by up to a factor of 40. Anomalous diffusion and two-component Brownian diffusion models were added to broaden the scope of biophysically-relevant datasets that can be analyzed using fcsSOFI. fcsSOFI was also packaged into a user-friendly GUI which can easily be outfitted with new diffusion models by the user. The application of fcsSOFI to simulations of fibrinogen diffusing in various matrix densities of polyacrylamide revealed more densely packed pores increased subdiffusive effects, while demonstrating the expanded biophysical scope of fcsSOFI. We hope that the updates in speed, scope, and usability allow a more diverse group of researchers to incorporate fcsSOFI into their research in topics like the extracellular matrix [5,47,48], chromatographic media [4,49], biofilms [50], cell membranes and cytosol [51,52], catalysis [53,54], and polymers [55].

## Appendix A fcsSOFI accurately characterizes anomalous diffusion dynamics

Datasets of diffusion at *D* = 1 × 10^5^ *nm*^2^*s*^−1^ (slower diffusion and simulation code modifications made to minimize confinement effects) with variable anomalous factors were generated to demonstrate the ability of fcsSOFI to accurately extract α. All averaged D and α values were comfortably within error of the values used to simulate the datasets.

## Appendix B Limited simulation complexity

Simulated datasets of fibrinogen diffusion were created as a simple case to demonstrate the efficacy of fcsSOFI as a super-resolution tool. Thus, it does not account for protein structure (each fibrinogen molecule is treated as a sphere), fibrinogen-fibrinogen and fibrinogen-matrix interactions, or the potential dynamic nature of the matrix. Experimental data of immobilized fibrinogen molecules was used to select experimentally plausible values for signal-to-noise ratio (SNR).

## Appendix C Experimental fibrinogen sample preparation

Alexa Fluor 546 labeled fibrinogen was purchased from Thermo Fisher Scientific and used without further modification. Fibrinogen is listed by the vendor as having 15 dyes per protein. Fibrinogen was diluted with 1x phosphate buffered saline (Thermo Fisher Scientific) to a 0.1 nM concentration. Rectangular number 1.5 glass coverslips (Corning, 22 x 30 mm2 size, 0.16-0.19mm thickness) were cleaned by first submerging the coverslips in ultrapure water (>18 mΩ), followed by a TL1 solution (6:1:1 (v/v) ratio of H_2_O:H_2_O_2_:NH_4_OH) at 75 °C for 90 s, followed by rinsing the slides once more in ultrapure water. The coverslips were then dried with ultra-high purity N_2_ (Airgas) before being treated in an O_2_ (Airgas) plasma cleaner (Harrick Plasma) at 11 W for 120 s. The fibrinogen solution was then drop-casted onto the cleaned glass cover slips and placed in a desiccator and protected from light overnight to dry. The sample was imaged the following day on a total internal reflection fluorescence (TIRF) microscope setup (Olympus IX83 body) equipped with a 561 nm diode laser (MGL-FN-561-100 mW, CNI). The sample was excited through a 100x, 1.49 NA objective (Olympus, UAPON-100XTIRF) at 2.0 mW/mm^2^ intensity. Emission from the sample was collected back through the objective and was separated from the excitation light with a dichroic mirror (DI03-R405/488/561/635-T1-25X36, Semrock) and notch filter (ZET561NF, Chroma Technology). Data was collected on a sCMOS camera (Photometrics Prime 95B) using a 50 ms integration time. Videos were collected at three different locations on each sample, and protein intensity values were retrieved using an established super-resolution localization program [56] from the first frames alone in order to mitigate the effects of photobleaching.

## Acknowledgements

The authors thank the Case Western Reserve University College of Arts and Sciences for financial support. S.Y. thanks support through a summer research scholarship provided by Case Western Reserve University SOURCE with additional funding provided by the Bruce Rakay Summer Research Fellowship.

## References

1. C. Dutta, L. D. C. Bishop, J. Zepeda O, S. Chatterjee, C. Flatebo, and C. F. Landes, “Imaging switchable protein interactions with an active porous polymer support,” J. Phys. Chem. B 124(22), 4412–4420 (2020).

2. B. Stempfle, A. Große, B. Ferse, K. F. Arndt, and D. Wöll, “Anomalous diffusion in thermoresponsive polymer-clay composite hydrogels probed by wide-field fluorescence microscopy,” Langmuir 30(46), 14056–14061 (2014).

3. F. Xiao, J. Hrabe, and S. Hrabetova, “Anomalous extracellular diffusion in rat cerebellum,” Biophys. J. 108(9), 2384–2395 (2015).

4. W. Calabrase, L. D. C. Bishop, C. Dutta, A. Misiura, C. F. Landes, and L. Kisley, “Transforming separation science with single-molecule methods,” Anal. Chem. 92(20), 13622–13629 (2020).

5. N. K. Reitan, A. Juthajan, T. Lindmo, and C. de Lange Davies, “Macromolecular diffusion in the extracellular matrix measured by fluorescence correlation spectroscopy,” J. Biomed. Opt. 13(5), 054040 (2008).

6. B. M. Regner, D. Vučinić, C. Domnisoru, T. M. Bartol, M. W. Hetzer, D. M. Tartakovsky, and T. J. Sejnowski, “Anomalous diffusion of single particles in cytoplasm,” Biophys. J. 104(8), 1652–1660 (2013).

7. N. Gröner, J. Capoulade, C. Cremer, and M. Wachsmuth, “Measuring and imaging diffusion with multiple scan speed image correlation spectroscopy,” Opt. Express 18(20), 21225 (2010).

8. D. J. Mai and C. M. Schroeder, “100th anniversary of macromolecular science viewpoint: single-molecule studies of synthetic polymers,” ACS Macro Lett. 9(9), 1332–1341 (2020).

9. L. M. Joubert, “Visualization of hydrogels with variable-pressure sem,” Microsc. Microanal. 15(SUPPL. 2), 1308–1309 (2009).

10. M. Maaloum, N. Pernodet, and B. Tinland, “Agarose gel structure using atomic force microscopy: gel concentration and ionic strength effects,” Electrophoresis 19(10), 1606–1610 (1998).

11. D. Axelrod, D. E. Koppel, J. Schlessinger, E. Elson, and W. W. Webb, “Mobility measurement by analysis of fluorescence photobleaching recovery kinetics,” Biophys. J. 16(9), 1055–1069 (1976).

12. A. B. Houtsmuller, “Fluorescence recovery after photobleaching: application to nuclear proteins,” Adv. Biochem. Eng. Biotechnol. 95177–199 (2005).

13. G. Baumann, I. Gryczynski, and Z. Földes-Papp, “Anomalous behavior in length distributions of 3d random brownian walks and measured photon count rates within observation volumes of single-molecule trajectories in fluorescence fluctuation microscopy,” Opt. Express 18(17), 17883 (2010).

14. A. J. Berglund and H. Mabuchi, “Tracking-fcs: fluorescence correlation spectroscopy of individual particles,” Opt. Express 13(20), 8069 (2005).

15. K. Hassler, M. Leutenegger, P. Rigler, R. Rao, R. Rigler, M. Gösch, and T. Lasser, “Total internal reflection fluorescence correlation spectroscopy (tir-fcs) with low background and high count-rate per molecule,” Opt. Express 13(19), 7415 (2005).

16. N. Bag and T. Wohland, “Imaging fluorescence fluctuation spectroscopy: new tools for quantitative bioimaging,” Annu. Rev. Phys. Chem. 65(1), 225–248 (2014).

17. T. Wohland, X. Shi, J. Sankaran, and E. H. K. Stelzer, “Single plane illumination fluorescence correlation spectroscopy (spim-fcs) probes inhomogeneous three-dimensional environments,” Opt. Express 18(10), 10627 (2010).

18. Y. Tian, M. M. Martinez, and D. Pappas, “Fluorescence correlation spectroscopy: a review of biochemical and microfluidic applications,” Appl. Spectrosc. 65(4), 115 (2011).

19. T. Dertinger, R. Colyer, R. Vogel, M. Heilemann, M. Sauer, J. Enderlein, and S. Weiss, “Super-resolution Optical Fluctuation Imaging (SOFI) BT - Nano-Biotechnology for Biomedical and Diagnostic Research,” in E. Zahavy, A. Ordentlich, S. Yitzhaki, and A. Shafferman, eds. (Springer Netherlands, 2012), pp. 17–21.

20. T. Dertinger, R. Colyera, G. Iyer, S. Weiss, and J. Enderlein, “Fast, background-free, 3d super-resolution optical fluctuation imaging (sofi),” Proc. Natl. Acad. Sci. U. S. A. 106(52), 22287–22292 (2009).

21. L. Kisley, R. Brunetti, L. J. Tauzin, B. Shuang, X. Yi, A. W. Kirkeminde, D. A. Higgins, S. Weiss, and C. F. Landes, “Characterization of porous materials by fluorescence correlation spectroscopy super-resolution optical fluctuation imaging,” ACS Nano 9(9), 9158–9166 (2015).

22. E. L. Elson and D. Magde, “Fluorescence correlation spectroscopy. i. conceptual basis and theory,” Biopolymers 13(1), 1–27 (1974).

23. P. Schwille and J. Ries, “Principles and Applications of Fluorescence Correlation Spectroscopy (FCS),” in (Springer, Dordrecht, 2011), pp. 63–85.

24. P. Schwille, J. Korlach, and W. W. Webb, “Fluorescence correlation spectroscopy with single□molecule sensitivity on cell and model membranes,” Cytometry 36(3), 176–182 (1999).

25. D. Wöll, “Fluorescence correlation spectroscopy in polymer science,” RSC Adv. 4(5), 2447–2465 (2014).

26. A. Przybylski, B. Thiel, J. Keller-Findeisen, B. Stock, and M. Bates, “Gpufit: an open-source toolkit for gpu-accelerated curve fitting,” Sci. Rep. 7(1), 1–9 (2017).

27. M. Shayegan, R. Tahvildari, K. Metera, L. Kisley, S. W. Michnick, and S. R. Leslie, “Probing inhomogeneous diffusion in the microenvironments of phase-separated polymers under confinement,” J. Am. Chem. Soc. 141(19), 7751–7757 (2019).

28. D. Matthews, “Supercharge your data wrangling with a graphics card,” Nature 562(7725), 151–152 (2018).

29. The New Category In SCMOS Cameras 10 Megapixel 6.5 Mm Pixel Size 500 Frames Per Second 29.4 Mm Field Of View 95% Quantum Efficiency (n.d.).

30. R. Haase, L. A. Royer, P. Steinbach, D. Schmidt, A. Dibrov, U. Schmidt, M. Weigert, N. Maghelli, P. Tomancak, F. Jug, and E. W. Myers, “CLIJ: gpu-accelerated image processing for everyone,” Nat. Methods (n.d.).

31. T. Quan, P. Li, F. Long, S. Zeng, Q. Luo, P. N. Hedde, G. U. Nienhaus, and Z.-L. Huang, “Ultra-fast, high-precision image analysis for localization-based super resolution microscopy,” Opt. Express 18(11), 11867 (2010).

32. H. Kang, S.-W. Lee, E.-S. Lee, S.-H. Kim, and T. G. Lee, “Real-time gpu-accelerated processing and volumetric display for wide-field laser-scanning optical-resolution photoacoustic microscopy,” Biomed. Opt. Express 6(12), 4650 (2015).

33. I. K. Park, N. Singhal, M. H. Lee, S. Cho, and C. Kim, “Design and performance evaluation of image processing algorithms on gpus,” IEEE Trans. Parallel Distrib. Syst. 23(1), 91–104 (2011).

34. J. T. Cooper and J. M. Harris, “Imaging fluorescence-correlation spectroscopy for measuring fast surface diffusion at liquid/solid interfaces,” Anal. Chem. 86(15), 7618–7626 (2014).

35. N. Fatin-Rouge, K. Starchev, and J. Buffle, “Size effects on diffusion processes within agarose gels,” Biophys. J. 86(5), 2710–2719 (2004).

36. T. Kihara, J. Ito, and J. Miyake, “Measurement of biomolecular diffusion in extracellular matrix condensed by fibroblasts using fluorescence correlation spectroscopy,” PLoS One 8(11), e82382 (2013).

37. D. Boening, T. W. Groemer, and J. Klingauf, “Applicability of an em-ccd for spatially resolved tir-ics,” Opt. Express 18(13), 13516 (2010).

38. J. Li and D. J. Mooney, “Designing hydrogels for controlled drug delivery,” Nat. Rev. Mater. 1(12), (2016).

39. Y. Qiu and K. Park, “Environment-sensitive hydrogels for drug delivery,” Adv. Drug Deliv. Rev. 53(3), 321–339 (2001).

40. A. Saini and L. Kisley, “Fluorescence microscopy of biophysical protein dynamics in nanoporous hydrogels,” J. Appl. Phys. 126(8), 81101 (2019).

41. L. Kisley, K. A. Serrano, D. Guin, X. Kong, M. Gruebele, and D. E. Leckband, “Direct imaging of protein stability and folding kinetics in hydrogels,” ACS Appl. Mater. Interfaces 9(26), 21606–21617 (2017).

42. L. M. Lira, K. A. Martins, and S. I. C. de Torresi, “Structural parameters of polyacrylamide hydrogels obtained by the equilibrium swelling theory,” Eur. Polym. J. 45(4), 1232–1238 (2009).

43. J. R. Tse and A. J. Engler, “Preparation of hydrogel substrates with tunable mechanical properties,” Curr. Protoc. Cell Biol. (SUPPL. 47), 10.16.1-10.16.16 (2010).

44. “» What are the time scales for diffusion in cells?,” http://book.bionumbers.org/what-are-the-time-scales-for-diffusion-in-cells/.

45. K. Chen, S. M. Anthony, and S. Granick, “Extending particle tracking capability with delaunay triangulation,” Langmuir 30(16), 4760–4766 (2014).

46. “KisleyLabAtCWRU (Kisley Research Group @ CWRU) · GitHub,” https://github.com/KisleyLabAtCWRU.

47. J. Tønnesen, V. V. G. K. Inavalli, and U. Valentin Nä Gerl, “Super-resolution imaging of the extracellular space in living brain tissue resource super-resolution imaging of the extracellular space in living brain tissue,” Cell 1721108–1111.e15 (2018).

48. C. Paviolo, F. N. Soria, J. S. Ferreira, A. Lee, L. Groc, E. Bezard, and L. Cognet, “Nanoscale exploration of the extracellular space in the live brain by combining single carbon nanotube tracking and super-resolution imaging analysis,” Methods 17491–99 (2020).

49. N. A. Moringo, H. Shen, L. D. C. Bishop, W. Wang, and C. F. Landes, “Enhancing analytical separations using super-resolution microscopy,” Annu. Rev. Phys. Chem. 69(1), 353–375 (2018).

50. J. M. Rodríguez-Suárez, C. S. Butler, A. Gershenson, and B. L. T. T. Lau, “Heterogeneous diffusion of polystyrene nanoparticles through an alginate matrix: the role of cross-linking and particle size,” 54(8), 5159–5166 (2020).

51. J. Sankaran and T. Wohland, “Fluorescence strategies for mapping cell membrane dynamics and structures,” APL Bioeng. 4(2), 020901 (2020).

52. R. F. Wissner, A. Steinauer, S. L. Knox, A. D. Thompson, and A. Schepartz, “Fluorescence correlation spectroscopy reveals efficient cytosolic delivery of protein cargo by cell-permeant miniature proteins,” ACS Cent. Sci. 4(10), 1379–1393 (2018).

53. J. Van Loon, A. V. Kubarev, and M. B. J. Roeffaers, “Correlating catalyst structure and activity at the nanoscale,” ChemNanoMat 4(1), 6–14 (2018).

54. J. Xie, J. Xu, X. Sun, H. Wang, D. A. Higgins, and K. L. Hohn, “Exploring microenvironment acidity inside the solvent-filled pores of mesoporous silica thin films via single-molecule spectroscopy,” J. Phys. Chem. C 123(33), 20333–20341 (2019).

55. M. Baier, D. Wöll, and S. Mecking, “Diffusion of molecular and macromolecular polyolefin probes in cylindrical block copolymer structures as observed by high temperature single molecule fluorescence microscopy,” Macromolecules 51(5), 1873–1884 (2018).

56. J. Chen, A. Bremauntz, L. Kisley, B. Shuang, and C. F. Landes, “Super-resolution mbpaint for optical localization of single-stranded dna,” ACS Appl. Mater. Interfaces 5(19), 9338–9343 (2013).

